# A Multi-objective Simulation-Optimization Framework for the Design of a Compliant Gravity Balancing Orthosis

**DOI:** 10.1101/2024.02.16.580745

**Authors:** Haider A. Chishty, Fabrizio Sergi

## Abstract

Flexion-synergy is a stereotypical movement pattern that inhibits independent joint control for those who have been affected by stroke; this abnormal co-activation of elbow flexors with shoulder abductors significantly reduces range of motion when reaching against gravity. While wearable orthoses based around compliant mechanisms have been shown to accurately compensate for the arm at the shoulder, it is unclear if accurate compensation can also be achieved while minimizing device bulk.

In this work, we present a novel, multi-objective simulation-optimization framework towards the goal of designing practical gravity-balancing orthoses for the upper-limb. Our framework includes a custom built VB.NET application to run nonlinear finite element simulations in SolidWorks, and interfaces with a MATLAB-based particle swarm optimizer modified for multiple objectives. The framework is able to identify a set of Pareto-optimal compliant mechanism designs, confirming that compensation accuracy and protrusion minimization are indeed conflicting design objectives.

The preliminary execution of the simulation-optimization framework demonstrates a capability of achieving designs that compensate for almost 90% of the arm’s gravity or that exhibit an average protrusion of less than 5% of the arm length, with different trade-offs between these two objectives.

## I. INTRODUCTION

**A**PPROXIMATELY 795 000 people experience new or recurrent strokes in the U.S. each year [1]; the resulting damage to the central nervous system (CNS) frequently results in the limitation of possible voluntary movements [2], [3]. The damaged CNS, upon exhausting all intact voluntary control pathways during functional task performance, will begin to rely on undamaged pathways that are non-specific to voluntary function [2]. This can result in the involuntary activation in distal limb flexors during shoulder motion, often termed “flexion synergy”, which can significantly inhibit independent joint control [2], [4], [5], [6], [7], [8], [9], [10]. A common flexion synergy, that between shoulder abductors and elbow flexors, reduces active range of motion (ROM) when reaching against gravity [2], [4], [5], [6], [7], [8], [9], [10]. However, ROM can be restored by relieving the shoulder muscles through compensation of the upper-limb’s weight, decoupling the flexion synergy [6], [7], [8], [10], [11], [12]. In clinical settings, this can be achieved using gravity-balancing technologies such as wearable robotics and exoskeletons [12], [13], [14], [15]; however, these devices typically possess inherent drawbacks associated with portability and load on the user, and therefore paradigms to decouple flexion synergies outside of controlled settings are far less common [7], [8].

While spring-based, passive devices, such as the occupational exoskeletons presented in [16], [17], [18], [19], [20], and [21], have been successful at providing wearable solutions to gravity compensation, such devices require rigid links to enforce spring extension [7], [8]. To avoid the use of rigid bodies, passive designs implementing compliant beams as flexure springs have been increasingly investigated due to their high customization [6], [7], [8]. Unfortunately, the relationship between the shape of a compliant beam and the compensation it provides is complex – therefore, optimization methods have been developed to determine ideal beam shapes that provide torques about the shoulder necessary to compensate for gravity. Combining finite element modeling (FEM) with shape optimization techniques, the groups in [6], [7] and [8] have been able to identify beam shapes that can compensate for gravitational torques at the elbow, shoulder, and both simultaneously, demonstrating the upper-limb can be balanced by more than 80% experimentally, and even more so in simulation.

However, in designing a compliant beam that perfectly compensates for gravity, it is possible that the compactness of the device is compromised. While the above groups impose penalties in their optimization cost functions relating to device size, the exact trade-off between compensation accuracy and device compactness has yet to be investigated. To investigate this trade-off, we have developed a novel, multi-objective simulation-optimization framework to determine compliant beam shapes capable of supporting the shoulder against the weight of the arm over all shoulder rotations in the sagittal plane: the framework modifies and extends single-objective particle-swarm optimization (PSO) to consider multiple objectives, using methods similar to those described in [22]. Using this framework, we are able to study how an ideal beam design changes when shifting priority away from compensation accuracy by examining pareto-optimal solutions.

## II. METHODS

### A. Problem Definition

The framework presented here aims to determine the shape of a compliant beam suitable to balance the weight of the human arm. The system is simplified to a single degree of freedom (DoF) by constraining motion to the sagittal plane and only considering cases when the elbow is fully extended, similar to [8] (Fig. 1A). The two ends of the compliant beam interface with the torso and arm respectively; at each interface, position and orientation are imposed relative to the body the end is attached to (Fig. 1B).

**Fig. 1.**
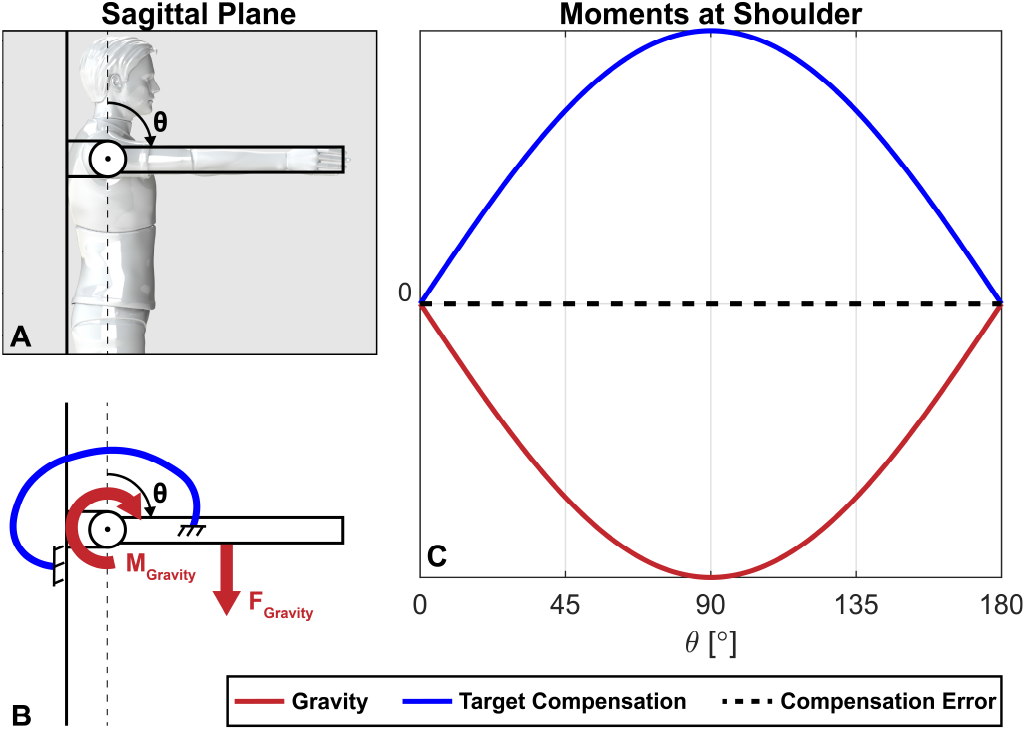
The system to be balanced is simplified by constraining rotation to the sagittal plane and enforcing the elbow to be fully extended (A). A compliant beam is used to compensate for gravity at the shoulder (B). Perfect compensation occurs when the reaction profile is sinusoidal (C).

The primary optimization objective of the framework is to compensate for the moment at the shoulder due to the weight of the upper-limb. This moment is a sinusoid function dependent on shoulder angle (Fig. 1C). Amplitude of the sinusoid is dependent on the arm’s mass. As such, to achieve perfect compensation of the arm’s weight, the reaction loading the compliant beam imposes on the arm must cause a sinusoid moment about the shoulder in the opposite direction to gravity.

The secondary objective of the framework is to identify designs that minimally protrude from the body; the implementation of this objective allows for the investigation of possible design objective trade-offs relevant to gravity compensation.

### B. Framework Overview

The purpose of this framework is to output the Pareto front of the optimization problem, enabling the study of trade-offs between gravity balancing capabilities and device protrusion. The framework consists of two components: 1) A finite element simulation performed in SolidWorks (Dassault Systèmes, Vélizy-Villacoublay, France); and 2) a particle swarm [23] optimization performed in MATLAB (MathWorks, Portola Valley, CA, United States) using the Global Optimization Toolbox, augmented to perform multiple-objective optimizations using methods similar to [22] (Fig. 2). Simulations and result exportation are automated using a custom application written in Visual Basic .NET (Microsoft, Redmond, WA, United States).

**Fig. 2.**
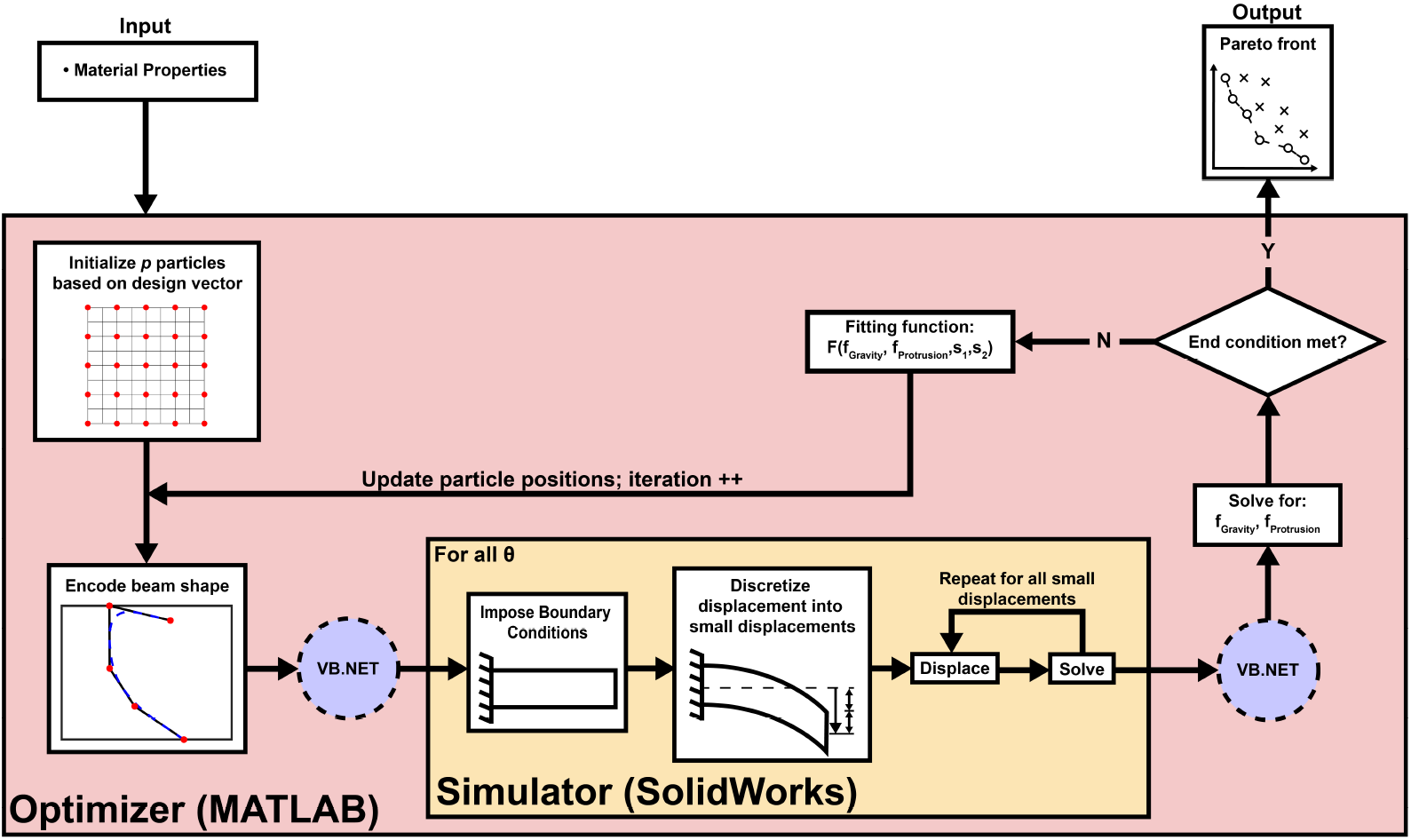
The multi-objective simulation-optimization framework. By implementing a SolidWorks finite element simulation into a particle-swarm optimizer in MATLAB, the framework will output the Pareto front of the optimization problem. A custom VB.NET application was written to communicate between MATLAB and SolidWorks.

### C. Shape Encoding

The design vector **x** (1) of the optimization problem is used to define the in-plane contour of a compliant beam (Fig. 3).

**Fig. 3.**
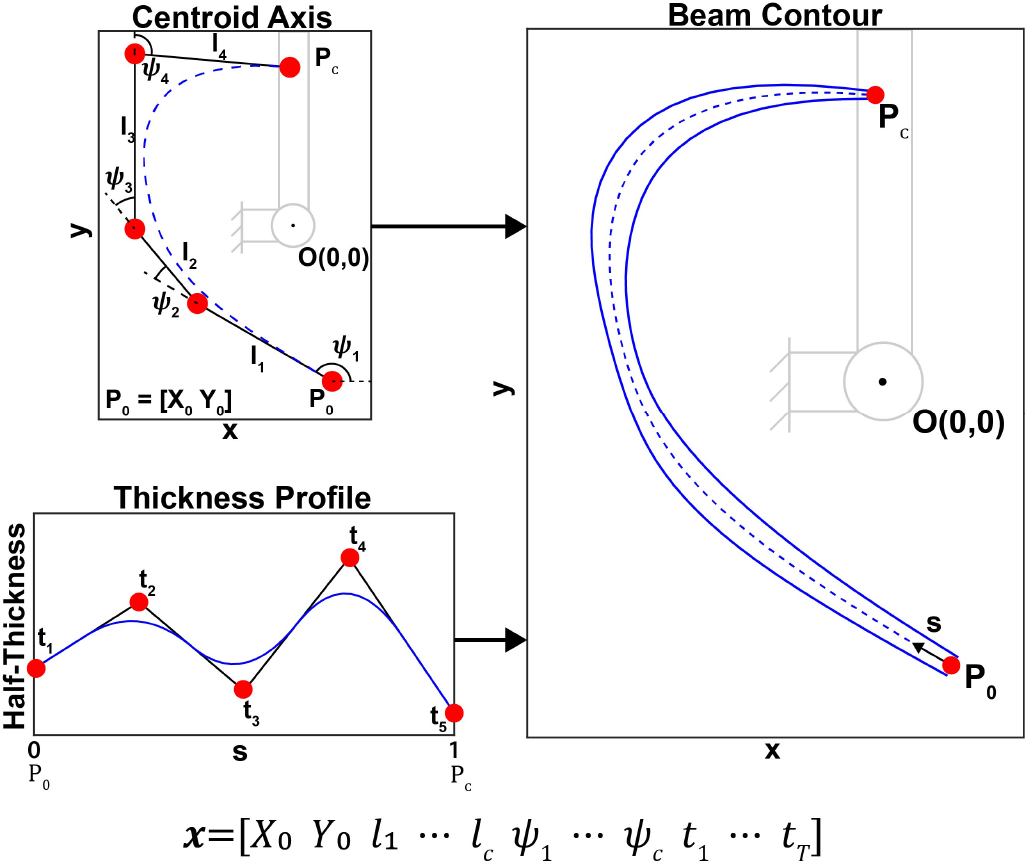
Shape encoding is performed in two stages: 1) a centroid axis is determined as a spline defined off a control polygon (Centroid Axis); 2) Bidirectional thickness is defined off the centroid axis by creating a thickness profile using methods similar to those used for the centroid axis (Thickness Profile).

Bounds on each design parameter can be seen in Table I.

**Table I.** Design Parameter Bounds

| Parameter | Units | Bounds |
| --- | --- | --- |
| $X_0$ | [m] | [-0.015 0.015] |
| $Y_0$ | [m] | [-0.3 0] |
| $l_{1:c}$ | [m] | [0.01 0.15] |
| $\psi_1$ | [rad] | $[-\pi \pi]$ |
| $\psi_{2:c}$ | [rad] | [-2 2] |
| $t_{1:T}$ | [m] | [0.005 0.02] |

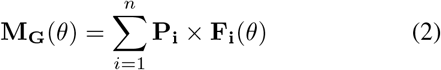

This definition is performed in MATLAB: the centroid axis of the beam is first defined off a control polygon using uniform subdivision [24]. The initial control point of the polygon is defined relative to the shoulder using parameters *X*_0_ and *Y*_0_. The subsequent *c* control points are defined using the next 2*c* design parameters; using linkage chain formulation [25], each subsequent control point is a distance *l*_*i*_ and relative angle *ψ*_*i*_ from the previous (Fig. 3: Centroid Axis). The initial control point *P*_0_ and final control point *P*_*C*_ correspond to the orthosis-torso and orthosis-arm interfaces respectively. For this study, *c* was set to 8. In the case of self-intersections, where the centroid axis would overlap with itself, the portion of the centroid axis beyond the intersection point is removed, and the intersection point is smoothed.

To define the surfaces of the compliant beam, an inplane thickness profile is created using the final *T* parameters *t*_*j*_; these parameters represent uniformly distributed y-coordinates used to define another spline through uniform subdivision; this spline represents the perpendicular thickness of the beam bidirectionally at any point on the centroid axis (Fig. 3: Thickness Profile). For this study *T* was set to 5.

### D. Simulation

A custom Visual Basic .NET application is used to import and extrude the contour of the compliant beam from MATLAB into SolidWorks and to automate the simulation. – the extrusion parameter is selected arbitrarily as it does not affect the shape of the reaction profile beyond linear scaling.

The 3D solid is deformed using a series of small load steps, allowing for the implementation of large deformation analysis. Results are calculated for each load step. Two boundary conditions are applied: 1) the orthosis-torso interface is kinematically fixed; 2) the orthosis-arm interface is rotated 180° clockwise about an axis coinciding with the location of the shoulder joint. A solid mesh comprised of linear, tetrahedral elements, with a maximum side length of 25 mm is created. Results from the simulation are then exported to be interpreted within MATLAB.

### E. Objective Calculations

For each load step of the simulation, the resulting reaction moment about the shoulder is calculated by summing the cross products of the position vectors of the nodes at one of the orthosis-body interfaces (in our case, the orthosis-torso interface) and their corresponding force vectors, (2):

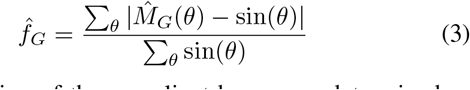

where **M**_G_(*θ*) is the reaction moment at the shoulder at any rotation *θ, n* is the number of nodes at the interface, *P*_*i*_ is the position vector of node *i* relative to the shoulder, and *F*_*i*_(*θ*) is the reaction force vector of node *i* at rotation *θ*.

Performing this calculation over each load step provides the reaction profile over the entire desired ROM. This profile is then normalized by its maximum value and compared to the target sinusoidal profile; the error between these functions is summed over different postures and normalized by the area of the sine wave. The resulting metric 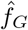 is the normalized gravity compensation error, and is the primary objective to be minimized by the optimizer (3).

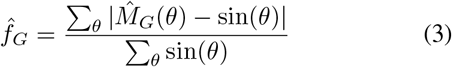

The protrusion of the compliant beam was determined as the average of the protrusions of all points on the beam, over all rotations. For a given point and rotation, protrusion is defined as the minimum distance between that point and the neutral axes of the torso or arm. The distance between the orthosis-arm interface and the neutral axis of the arm is also computed and subsequently added to the previous definition of protrusion to determine a protrusion index. This correction was introduced to penalize beam shapes that would require large components in order to attach to the arm.

The secondary objective of the optimizer, 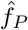, is calculated by normalizing the protrusion index by 0.6 m, scaling the protrusion to a convenient range. For this study, only the initial and final shoulder angles were used to calculate protrusion, rather than all rotations.

### F. Optimization

PSO was selected for use in the framework due to the large number of design parameters necessary to define the shape of a compliant beam (23 for this study) [23]. For a single objective optimization, PSO begins by initializing *p* particles, each corresponding to individual design vectors. For this study, *p* was set to 50. At every iteration, the objective function is evaluated based on each particle. These evaluations influence the evolution of each particle in the subsequent iteration. The update, or velocity, vector *V* of each particle is based on three components: a particle’s current trajectory, a particle’s best evaluation over all iterations, and the cumulative best evaluation over all particles and iterations. This update vector can be seen in (4):

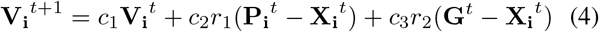

where *i* and *t* represent the current particle and iteration respectively, *X*_*i*_ is the position of particle *i, P*_*i*_ is the particle position at particle *i*’s best evaluation, and *G* is the particle position for the overall best evaluation. *c*_1_, *c*_2_, and *c*_3_ are weighting constants used to place emphasis on specific components of the update function; *r*_1_ and *r*_2_ are random coefficients between 0 and 1 updated every iteration.

To investigate trade-offs between multiple objectives, the single objective PSO must be augmented. Using methods similar to those described in [22], several modifications are made, as described below.

First, the evaluation for every particle across every iteration is logged in objective space - the multi-dimensional space described by the multiple design objectives; this allows each compliant beam design to be categorized into two groups: those that are dominated and those that are not. A solution is classified as dominated if there exists at least one other solution with: 1) equal or more optimal values in all components, and 2) a more optimal value in at least one component. The set of non-dominated solutions is the Pareto set, or Pareto frontier, of the multi-objective optimization problem, and demonstrates the trade-off between the problem’s multiple objectives. To ensure adequate exploration of the multi-dimensional objective space, specifically along the Pareto frontier, two further modifications are implemented.

The objective function *F* (**x**) of the multi-objective optimization problem is set to be the weighted sum of normalized objectives 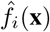 (5):

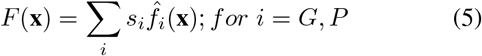

where *s*_*i*_ represents the weight of objective *i*. Each weight changes sinusoidally with iteration *t* with differing phase (6a, 6b). This weighting is used to promote exploration over the objective space, such that priority between objectives changes:

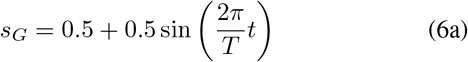

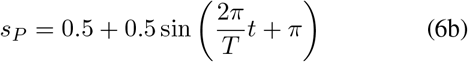

where for this study the period *T* was selected as 10 iterations, and phases of the two functions were offset by half a period (i.e., functions always summed to 1).

The global best solution of the optimization problem (*G* in (4)) is also modified in our framework: rather than corresponding to the solution that provides the best cost function over all particles, it is defined as the most isolated point along the Pareto front. Isolation is defined as the minimum distance from one Pareto solution to any other solution on the Pareto front (in objective space), such that the most isolated Pareto solution has the largest minimum distance.

In the event that a particle design vector leads to an unsuccessful evaluation, both normalized objectives 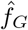 and 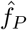 are assigned a value of 2. These cases can occur if beam self-intersection occurs during displacement, the VB.NET application fails to successfully communicate with either platform, or the simulation is unable to complete a full rotation (e.g. due to simulated buckling). For this study, the condition of a full rotation is softened to a minimum of 150°.

## III. RESULTS

The framework was run based on material properties corresponding to commercially available stereolithography 3D printing, compliant resin (elastic modulus *E* = 1.7 MPa, Poisson’s ratio *v* = 0.45, mass density *ρ* = 1250 kg/m^3^). The framework was run for a total of 30 iterations, allowing for three full period of objective weight oscillations.

### A. Framework Results

Of the 1500 design vectors evaluated, 471 were feasible solutions; 717 solutions were discarded due to insufficient rotation, and the remaining 312 were discarded due to intersections or application errors. All non-discarded design solutions can be seen in Fig. 4. As can be seen, the Pareto front is identified as the set of non-dominated solutions. These Pareto-optimal solutions offer good contrast between those that prioritize 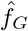 versus those that prioritize 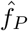, as well as those that prioritize both similarly. Three Pareto-optimal designs (labeled A:C in Fig. 4) are presented in the subsequent section.

**Fig. 4.**
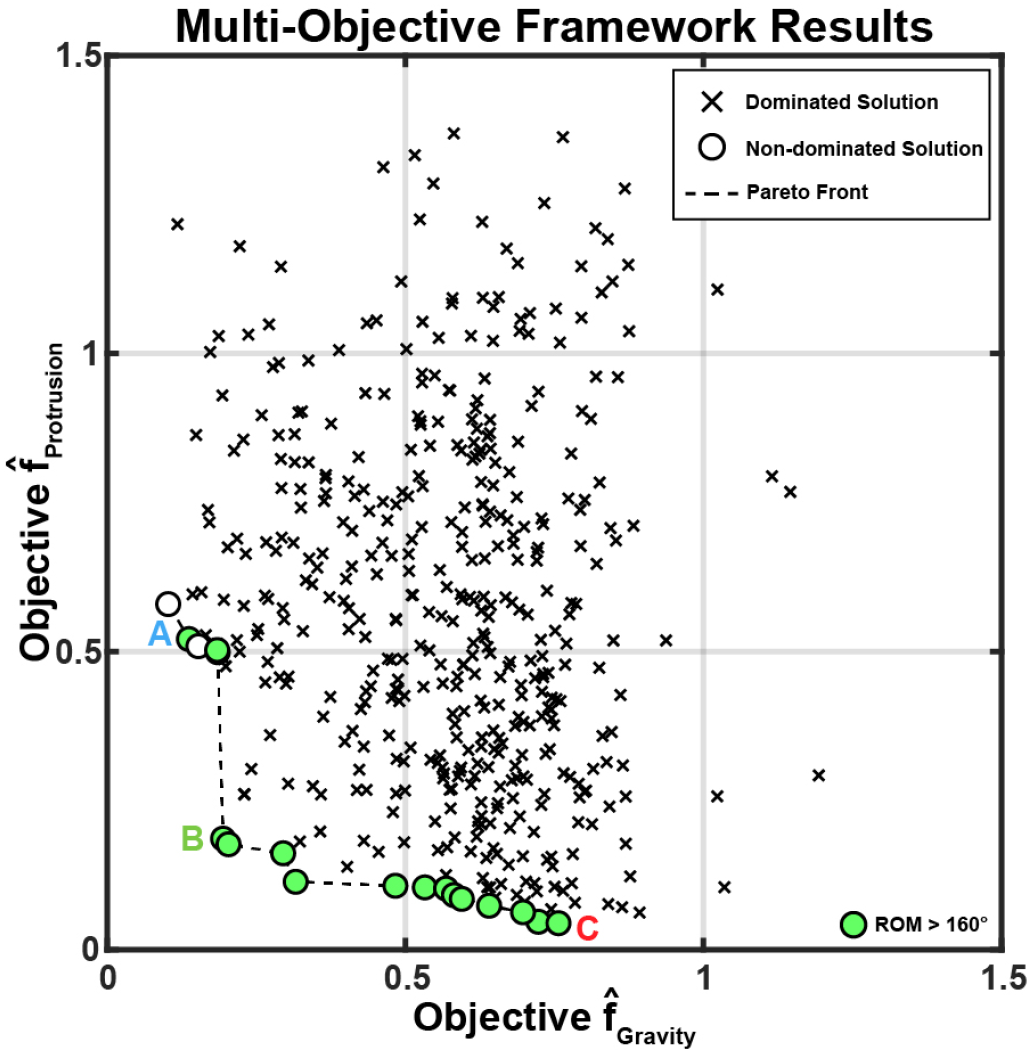
Multi-objective optimization results. The Pareto front is defined as the set of non-dominated solutions and represents the trade-off between the two design objectives. Pareto-optimal solutions capable of reaching 160° rotation are labelled.

### B. Pareto Optimal Designs

Three Pareto-optimal solutions outputted from the framework are seen in Fig. 5. The solutions were selected among those that could rotate at least 160°.

**Fig. 5.**
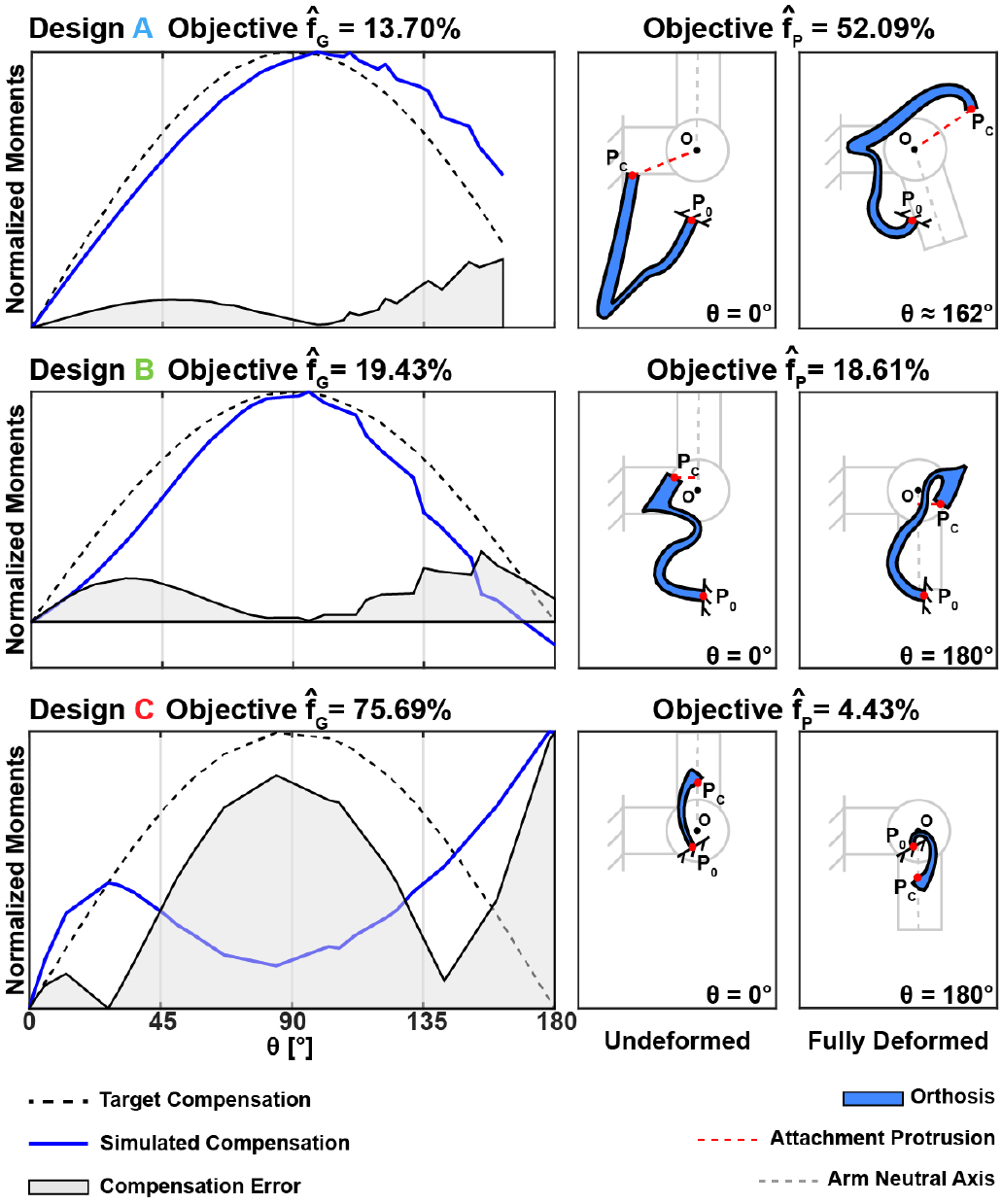
Pareto Optimal Solutions. Design A exhibits exceptional compensation accuracy but large protrusions; Design B exhibits acceptable levels of compensation and smaller protrusions; Design C exhibits poor compensation accuracy but minimal protrusions.

Design A offers a sinusoid moment profile (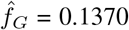) indicating that this design can very accurately compensate for the weight of the arm. However, the design shows infeasibility as, in addition to a region of extreme protrusion, the attachment point *P*_*C*_ is far off the arm, requiring an additional mechanism to ensure correct kinematic constraints (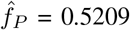).

In contrast, Design C offers exceptionally low protrusion (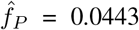) while resulting in a compensation profile that would be unusable for any practical gravity-balancing application (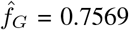).

Design B provides suitable compromise: relative to Design A, the solution offers only 6% less compensation (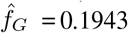) while drastically reducing its overall protrusion; further, the attachment point *P*_*C*_ is very close to the arm’s neutral axis (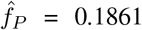). Such a solution would be suggested as a suitable design to be fabricated as a wearable prototype.

## IV. DISCUSSION AND CONCLUSION

### A. Discussion

Towards the goal of designing effective and feasible orthoses for the upper-limb, a novel, multi-objective simulation-optimization framework is presented. This framework enables the study of trade-offs between different design objectives and can successfully identify a set of Pareto-optimal compliant beam designs.

As can be seen from Fig. 4, the presence of a Pareto front indicates that the two design criteria are indeed conflicting, and that trade-off is necessary if one objective is to be prioritized. This aligns with the expectation: seeking to minimize device protrusion excludes a subset of designs that are capable of providing accurate compensation.

Three Pareto-optimal designs are presented. As described above, Design A prioritizes compensation accuracy, but offers low feasibility due to the arm attachment being far removed; alternatively Design C prioritizes a low-profile design while providing inadmissible amounts of gravity compensation; Designs B offer adequate compromise between the two design objectives by exhibiting a sinusoid compensation profile while also offering acceptable amounts of protrusion.

While Designs A exhibits extreme protrusions, its compensation capabilities align with experimental results presented in [7] and [8], where errors in gravity compensation of the elbow and shoulder (both presented as 1 DoF systems) are less than 20%. Simulated compensation presented in [7] can be as low as 2%, however, indicating that the framework presented here may require additional optimization iterations to find such a solution, or that such extreme compensation is only possible by compromising the minimization of protrusion.

Orthoses developed from this framework or those similar offer uses in both assistive and rehabilitative spaces: with accurate compensation, shoulder strength is no longer needed and shoulder abduction/adduction ROM is returned; further, as shoulder muscles are alleviated, distal limb flexion synergy is decoupled, returning reaching ROM as well. However, full compensation is not desirable if flexion synergy is to be rehabilitated as studies have shown that reaching under full shoulder support does not improve reaching ROM over conventional methods [10], [11], [26]; instead, shoulder abduction support should progressively be decreased [10], [11], [26]. In such cases, designs outputted from this framework can be scaled down to undercompensate the shoulder by a controllable amount. This application could potentially be augmented by fabricating the orthoses from materials with tunable stiffnesses [27], [28].

### B. Future Work and Conclusion

The primary goal of future iterations of the framework should be to increase computational efficiency by eliminating errors associated with insufficient ROM. This can be done by validating simulation results using different mesh densities and element classes, simulation types, or solver software. Results can also be validated through experimental protocols using fabricated prototypes.

To extend the effectiveness of designs found by the framework, the single DoF simplification can be lifted, extending the compensation target profile to be a function of both shoulder and elbow angle.

Finally, the particle swarm optimizer may be further modified to better promote exploration: this can be done by assigning particle-specific objective weights, such that each particle possesses a unique combination of priority. Further, alternate definitions of the global best solution could be explored, such as defining isolation in design vector space rather than objective space [22].

